# TITER: predicting translation initiation sites by deep learning

**DOI:** 10.1101/103374

**Authors:** Sai Zhang, Hailin Hu, Tao Jiang, Lei Zhang, Jianyang Zeng

## Abstract

**Motivation:** Translation initiation is a key step in the regulation of gene expression. In addition to the annotated translation initiation sites (TISs), the translation process may also start at multiple alternative TISs (including both AUG and non-AUG codons), which makes it challenging to predict TISs and study the underlying regulatory mechanisms. Meanwhile, the advent of several high-throughput sequencing techniques for profiling initiating ribosomes at single-nucleotide resolution, e.g., GTI-seq and QTI-seq, provides abundant data for systematically studying the general principles of translation initiation and the development of computational method for TIS identification.

**Methods:** We have developed a deep learning based framework, named TITER, for accurately predicting TISs on a genome-wide scale based on QTI-seq data. TITER extracts the sequence features of translation initiation from the surrounding sequence contexts of TISs using a hybrid neural network and further integrates the prior preference of TIS codon composition into a unified prediction framework.

**Results:** Extensive tests demonstrated that TITER can greatly outperform the state-of-the-art prediction methods in identifying TISs. In addition, TITER was able to identify important sequence signatures for individual types of TIS codons, including a Kozak-sequence-like motif for AUG start codon. Furthermore, the TITER prediction score can be related to the strength of translation initiation in various biological scenarios, including the repressive effect of the upstream open reading frames (uORFs) on gene expression and the mutational effects influencing translation initiation efficiency.

**Availability:** TITER is available as an open-source software and can be downloaded from https://github.com/zhangsaithu/titer

## 1 Introduction

Translation initiation plays an important role in mRNA translation, in which the methionyl tRNA unique for initiation (Met-tRNAi) identifies the AUG start codon and triggers the downstream translation process (Jackson *et al.*, 2010; Sonenberg and Hinnebusch, 2009; Hershey *et al.*, 2012). As translation initiation is an essential step in controlling gene expression and protein synthesis, the dysregulation of the initiation process can cause various human diseases, including cancers and metabolic disorders (Sonenberg and Hinnebusch, 2009; Hershey *et al.*, 2012). On the other hand, the mechanisms underlying translation initiation, e.g., the recognition of a translation initiation site (TIS) by the 80S ribosome assembly, are far more complicated than scientists had initially believed (Gao *et al.*, 2015; Lee *et al.*, 2012). In particular, experimental studies have shown that the eukaryotic translation is not always initiated at the canonical AUG start codons (Gao *et al.*, 2015; Peabody, 1989; Lee *et al.*, 2012; Kozak, 1989). In addition to those annotated translation initiation sites (aTISs), both upstream (uTISs) and downstream translation initiation sites (dTISs) can also occur at non-AUG codons, which yields alternative open reading frames (ORFs) that can be translated into short peptides or affect the expression levels of the main ORFs (Calvo *et al.*, 2009; Sonenberg and Hinnebusch, 2009; Jackson *et al.*, 2010; Hershey *et al.*, 2012; Barbosa *et al.*, 2013). This underscores the necessity of a better understanding of the mechanism of TIS recognition.

Ribosome profiling (ribo-seq), a high-throughput deep sequencing based technique that measures the ribosome protected fragments *in vivo*, has become a widely-used method to quantify the translation dynamics on a transcriptome (Ingolia *et al.*, 2009, 2012). However, the standard ribo-seq is not suitable for directly detecting TISs (Lee *et al.*, 2012). Based on ribo-seq, additional techniques, e.g., the global translation initiation sequencing (GTI-seq) (Lee *et al.*, 2012) and the quantitative translation initiation sequencing (QTI-seq) (Gao *et al.*, 2015), have been developed for a systematic mapping of the start codon positions at single-nucleotide resolution *in vivo*. By far, these techniques have provided abundant data for investigating the principles of translation initiation and translational control.

A variety of computational methods have been developed to predict alternative TISs or ORFs (Zien *et al.*, 2000; Hatzigeorgiou, 2002; Li and Jiang, 2005; Zur and Tuller, 2013; Chew *et al.*, 2016). However, most of these methods only focused on the AUG start codon and did not consider the widely-observed non-AUG TISs. In addition, few studies have utilized experimental data generated by GTI-seq or QTI-seq in their works, which limits the empirical predictive power of these methods. Recently, a linear regression based approach, called PreTIS, was proposed to predict non-canonical TISs by incorporating both AUG and its near-cognate codons (i.e., the codons differing from AUG by one nucleotide), in which TISs identified by GTI-seq were used to train the prediction model (Reuter *et al.*, 2016). However, only alternative TISs in the 5’ UTR (i.e., uTISs) were considered by PreTIS, and several constraints on the candidate TISs (e.g., the codon position in the transcript and the existence of an orthologous mouse sequence) had to be imposed due to the limitations of their feature engineering. To our best knowledge, our work is the *first* attempt to predict all possible TISs, including uTISs, aTISs and dTISs, in a unified framework without any restriction to the codon site of interest.

Recently, deep learning has become one of the most effective and powerful prediction methods in machine learning (Hinton and Salakhutdinov, 2006; Hinton *et al.*, 2006). It has been widely used and shown to be able to achieve the state-of-the-art prediction performance on various machine learning tasks, such as speech recognition (Hinton *et al.*, 2012), image classification (Hinton and Salakhutdinov, 2006) and natural language processing (Collobert *et al.*, 2011). In addition, deep learning is gradually gaining its popularity in bioinformatics and has yielded superior performance over conventional learning methods on a variety of biological prediction tasks, such as the predictions of protein-nucleotide binding (Zhang *et al.*, 2015; Alipanahi *et al.*, 2015), functional effects of noncoding sequence variants (Quang and Xie, 2016; Zhou and Troyanskaya, 2015) and ribosome stalling (Zhang *et al.*, 2016).

In this study, we have developed a deep learning based framework, named TITER (Translation Initiation siTE detectoR), for accurately predicting TISs based on the available high-throughput sequencing data. TITER possesses more flexibility than previous methods, and integrates the prior preference of the TIS codon composition as well as their surrounding sequence contexts into a unified framework, in which an ensemble of hybrid deep convolutional and recurrent neural networks is implemented to effectively and robustly capture the sequence features of translation initiation. Extensive validation tests have shown that TITER can greatly outperform the state-of-the-art computational approaches in detecting TISs. In addition, TITER can successfully identify significant sequence motifs for different TIS codons, including a Kozak-sequence-like motif for the AUG TIS codon. By combining gene expression data with TITER analysis, we found that the predictions of TITER well correlated with translation efficiency (TE) of genes, which basically reconfirmed the repressive effect of uORFs on gene expression. Furthermore, our comparative analyses on several important mutations around TISs showed that the fold changes of TITER prediction scores well conformed to the experimentally-verified mutational effects reported in the literature (Noderer *et al.*, 2014; Calvo *et al.*, 2009). These results demonstrated that TITER can offer a powerful tool to model the sequence features of translation initiation and identify potential TISs, which will provide useful insights into understanding the underlying mechanisms of translation initiation.

## 2 Methods

### 2.1 Datasets

We mainly used the QTI-seq data collected from the HEK293 cell line (Gao *et al.*, 2015) and the annotated TISs retrieved from Ensembl v84 (Aken *et al.*, 2016) (together denoted by Gao15) to train and test TITER. Specifically, we first retrieved all transcripts, each containing at least one translation initiation site identified by the QTI-seq experiment. Then we combined the QTI-seq identified TISs among these transcripts with the corresponding annotated TISs obtained from the Ensembl database, and regarded them as positive samples. To construct a dataset of negative samples that well reflected the imbalance of positive and negative samples *in vivo*, for each TIS in the positive dataset, we chose up to ten codon sites of the same triplet within the same transcript as negative samples. Considering the leaky scanning nature of the translation initiation process (Michel *et al.*, 2014), we searched for the negative samples starting from the 5’ end of an mRNA, until ten eligible sites were found. Altogether, the Gao15 dataset consisted of 9,776 positive samples and 94,899 negative samples from 4,111 transcripts, among which 400 transcripts were reserved for testing and the others were regarded as our training data (denoted by Gao15_test and Gao15_train, respectively).

To further evaluate the prediction performance of TITER in a scenario with an unlimited ratio of positive and negative samples, from the 400 test transcripts, we also constructed another test dataset (denoted by Gao15_test_extended), in which we identified all possible initiation codon sites before the last TIS of each transcript when searching for negative samples. Notably, as QTI-seq identified versatile TISs in terms of both codon composition and positions, our constructed training and test datasets also reflected this versatility, which thus enabled TITER to capture diverse sequence features of translation initiation that exist *in vivo*. In particular, we found that in the QTI-seq data derived from (Gao *et al.*, 2015), besides the canonical aTISs and the conventional AUG TISs, there also exist a large number of alternative TISs (i.e., uTISs and dTISs) and non-AUG TISs (e.g., CUG, GUG and UUG) (Fig. 2). This observation underscores the necessity of modeling the universal rules of translation initiation.

### 2.2 Modeling the preference of codon composition of TISs

It has been experimentally validated that there is a significant preference of codon composition among TISs (Lee *et al.*, 2012; Gao *et al.*, 2015). For example, the AUG codon generally plays a dominant role (>50%) at the TISs (Fig. 2(a)). Interestingly, other codons that differ from AUG at the first nucleotide (nt), including CUG, GUG and UUG, also greatly contribute to translation initiation (*∼*25%; Fig. 2(a)). Here, we collectively denote these four codons by NUG, in which “N” stands for any possible nucleotide. With this prior knowledge on codon preference, we propose to explicitly model the preference of codon composition of TISs by

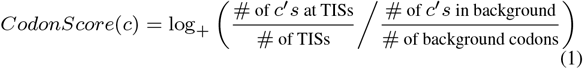

in which *CodonScore*(*c*) is defined as the preference of codon composition for a certain codon *c*, # denotes the number of the entities in the defined category and the background is defined as the upstream 1/3 segment of the transcripts, following the same principle as in (Gao *et al.*, 2015). In particular, we denote the function between the parentheses on the right-hand side of Eq. 1 by *f* (*c*). Then we define log_+_(*x*) = log(*x*) if log(*x*) > 0, log_+_(*x*) = *αx* otherwise, where *α* = min{log(*f* (*c*))| log(*f* (*c*)) > 0, *c* is a codon}. This definition is introduced as we notice that for those codons that are less similar with the canonical AUG start codon, their abundance in the QTI-seq dataset is actually lower than the background, leading to a negative *CodonScore* value if we only use the log(*x*) form. Although these codons only account for *∼*10% in the Gao15 dataset, to avoid the over-devaluation of the *CodonScores* for these codons due to the nature of logarithm function, we propose to use a linear model to define the *CodonScore* for these codons. In particular, the top five codons with the highest *CodonScores* value are AUG, CUG, GUG, UUG and ACG (Supplementary Fig. 1), which is consistent with the statistics shown in Fig. 2(a).

**Fig. 1.**
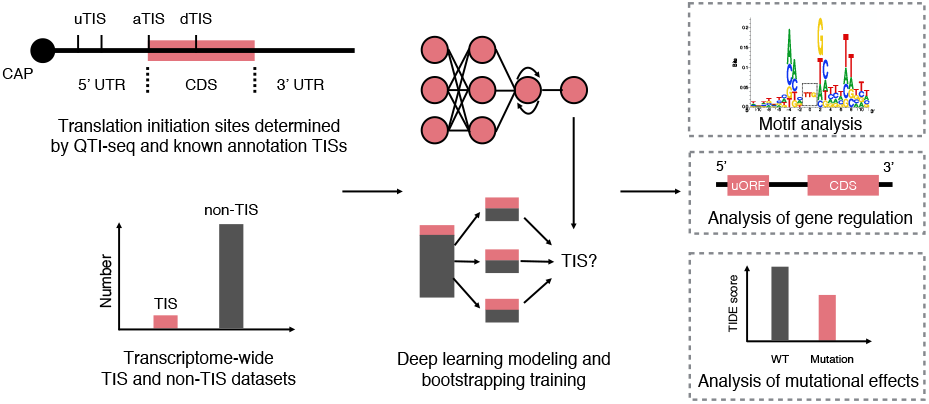
Schematic overview of the TITER pipeline. See the main text for more details.

**Fig. 2.**
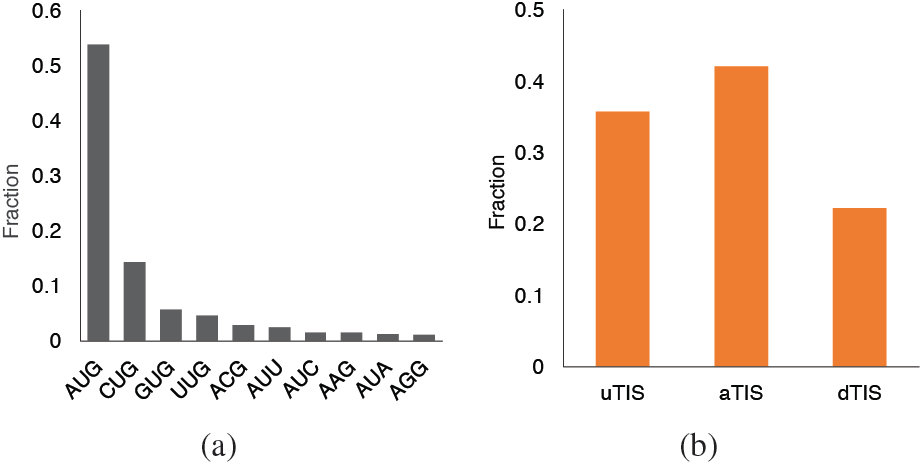
Statistics of the translation initiation sites in the Gao15 dataset. (a) Codon composition of TISs, in which only those codons with a fraction > 1% are shown. (b) Fractions of different types of TISs.

### 2.3 Modeling the contextual features of TISs

It has been indicated that the contextual sequences around the TISs can influence the likelihood of translation initiation (Kozak, 1989; Noderer *et al.*, 2014). For example, the upstream and downstream sequences of the canonical AUG TISs exhibit a consensus motif called the Kozak sequence (Kozak, 1989). This observation underlies the rationale of modeling the translation initiation events by encoding the contextual sequence features surrounding TISs.

Here, we develop a deep neural network to systematically model the sequence features of TISs (Fig. 3(a)). In particular, each TIS is extended both upstream and downstream by 100 nts, which yields the contextual profile of a translation initiation event and is denoted by *s* = (*n*_1_, *…, n*_203_), where *n*_*i*_ denotes the nucleotide at the *i*th position. To characterize the local motifs of the extended sequence *s*, we first encode the nucleotides using the one-hot encoding technique (Pedregosa *et al.*, 2011), that is, a nucleotide of a particular type (A, U, C, or G) is encoded by a binary vector of length four, in which the corresponding position is one while the others are zeros, after indexing all four types of nucleotides. Then we employ multiple convolution operators (denoted by conv(·)) to scan the encoded sequence profile and detect the local motifs around each TIS. After that, the pooling operators (denoted by pool(·)) are used to identify the activated motifs and also reduce the dimensions of hidden features. Note that the conventional convolution-pooling structures are order insensitive, as they only detect whether certain motifs exist regardless of their positions or orders. To further characterize the motif order that may also contribute to translation initiation, we also stack a long short-term memory (LSTM) network (denoted by LSTM(·)) upon the convolution-pooling module, which takes the pooled feature vectors as input and models the long-term dependencies between different motifs. Finally, the outputs of the LSTM at all positions are concatenated and fed into a logistic regression layer (denoted by logist(·)) to compute the probability of translation initiation for the input sequence. Indeed, when considering the LSTM outputs at all positions, we also implement the attention mechanism that has been widely used in deep learning (Denil *et al.*, 2012; Larochelle and Hinton, 2010) to leverage the relevance of each position to the final prediction. Altogether, by fully exploiting both sequence composition and motif orders, the hybrid convolutional and recurrent neural network in TITER computes the following score,

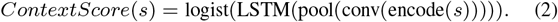

**Fig. 3.**
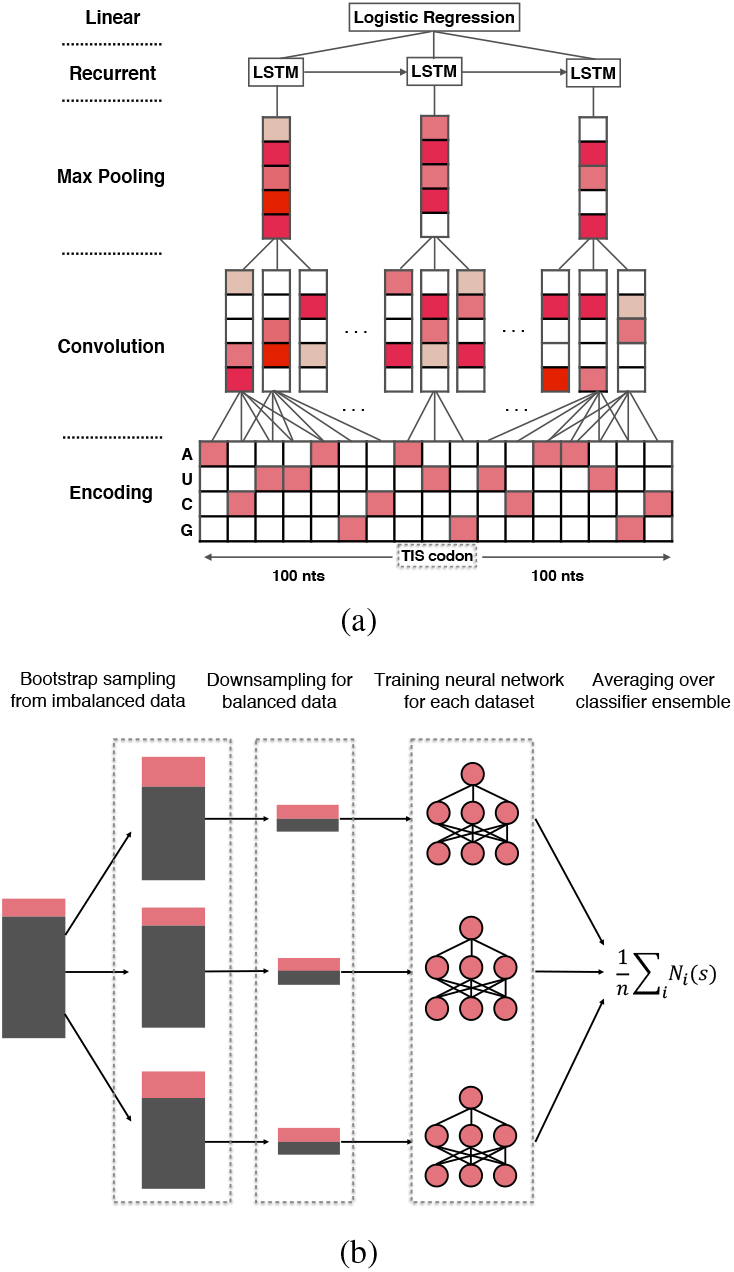
Schematic illustration of (a) the hybrid deep neural network architecture and (b) the bootstrapping-based technique used in TITER. See the main text for more details.

The sequence profiles of non-TISs (i.e., the negative samples) can be modeled in the same manner. Note that a similar hybrid neural network architecture has also been implemented in (Quang and Xie, 2016; Hassanzadeh and Wang, 2016) for different tasks, i.e., predicting the functional effects of noncoding mutations and the DNA binding protein targets, respectively.

### 2.4 Model training and model selection

Given the training samples {(*s*_*i*_, *y*_*i*_)}_*i*_, the loss function of our deep learning framework is defined as the sum of the negative log likelihoods (NLLs), i.e.,

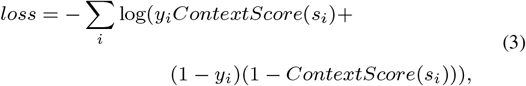

where *y*_*i*_ indicates whether *s*_*i*_ is the sequence profile of a TIS or not. We use the standard error backpropagation algorithm (Rumelhart *et al.*, 1986) and the batch gradient descent method (Bengio, 2012) to train the hybrid neural network and search for the network weights that minimize the loss function of Eq. 3. Several regularization techniques, including the max-norm constraints on weights (Srebro *et al.*, 2005), dropout (Srivastava *et al.*, 2014) and early stopping (Bengio, 2012), are also employed to optimize the training process and address the overfitting problem.

Our hybrid deep neural network architecture and various optimization techniques used in the training process have introduced several hyperparameters, e.g., the kernel size, kernel number and max-norm of weights, that need to be determined. A proper hyperparameter calibration procedure can help yield better solutions to the optimization problem in Eq. 3. Here, we use the tree-structured Parzen estimator (TPE) approach (Bergstra *et al.*, 2011) to calibrate the hyperparamters in our model, including kernel size, kernel number and max-norm of weights in the convolution layer, pooling length of the max-pooling layer, output dimension of the LSTM layer, dropout rate, and the optimizer algorithm. In particular, we first use all the positive training samples and an equal number of randomly-selected negative training samples to optimize the hyperparameters based on TPE (with 100 evaluations), and then choose the hyperparameters that achieve the minimum loss to further train our final models (Supplementary Table 1).

**Table 1.** Results on estimating translation efficiency (TE) from the TITER prediction scores, using a linear regression model with different combinations of features.

| Feature set | Feature coefficients in regression (mean±std) | Spearman’s correlation $r$ | $P$ value | MSE |
| --- | --- | --- | --- | --- |
| aTIS | $1.833 \pm 0.034$ | 0.234 | $1.077\text{e-}72$ | 3.081 |
| aTIS+uTIS | $1.774 \pm 0.040, -0.126 \pm 0.005$ | 0.245 | $3.801\text{e-}79$ | 3.063 |
| aTIS+AUG | $1.709 \pm 0.038, -0.029 \pm 0.005$ | 0.234 | $2.712\text{e-}72$ | 3.078 |
| aTIS+uTIS+lofUTR | $1.744 \pm 0.037, -0.122 \pm 0.005, -0.0001 \pm 3.634\text{e-}5$ | 0.236 | $2.228\text{e-}73$ | 3.063 |

“MSE” denotes the mean square error. “aTIS” represents the predicted TISScore for the aTIS, “uTIS” represents the sum of TISScores of all the eligible uTISs, “AUG” represents the number of AUGs in the upstream region of the aTIS, and “lofUTR” represents the length of the 5’ UTR. The mean and the standard deviation (std) of the feature coefficients in the regression model in a 10-fold cross-validation procedure were calculated.

We realize that the task of predicting TISs is an imbalanced classification problem (i.e., much more negative samples than positive ones), for which the standard training procedure designed for balanced samples cannot be appied directly. On the other hand, making use of more negative samples in the training process can lead to a more robust model with less variance in prediction (Wallace *et al.*, 2011). To tackle this imbalance problem, here we employ a bootstrapping-based technique (Fig. 3(b)) derived from the theory established in (Wallace *et al.*, 2011). Briefly, we first construct several groups of bootstrap samples (denoted by *S*_*i*_) from the original imbalanced population *S*, by randomly selecting samples with replacement. Then for each group *S*_*i*_, we balance the samples by downsampling, i.e., randomly selecting an equal number of positive and negative samples from *S*_*i*_, which yields a balanced dataset *B*_*i*_. After that, a hybrid deep neural network *N*_*i*_ is trained based on each dataset *B*_*i*_ independently, resulting in an ensemble of binary classifiers {*N*_*i*_}. Given an input sequence *s*, its final prediction score is averaged over the prediction scores output by all classifiers, i.e.,

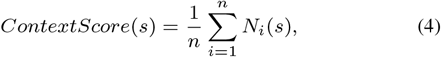

in which *n* is the total number of the constructed balanced datasets in {*B*_*i*_} (which is also equivalent to the total number of trained classifiers in {*N*_*i*_}). We apply this bootstrapping-based technique to our training dataset and train 32 independent deep neural networks. After that, their prediction scores are averaged and used as the final estimated probability of translation initiation for the given input sequence profile.

Due to the nature of non-convex optimization, random weight initialization can influence the search results of the gradient descent algorithm (Bengio, 2012). This initialization bias may also introduce variance to our modeling and further affect the prediction performance. In TITER, the aforementioned bootstrapping-based technique can alleviate such an initialization bias in addition to the sample bias, as the network weights have been initialized independently for each balanced sample group before the training process.

The hybrid deep neural network of TITER has been implemented using the Keras library^1^, and the Tesla K20c GPUs have been used to speed up the training process. The TPE algorithm for hyperparameter calibration has been implemented based on Hyperas^2^, a Python library for optimizing hyperparameters of the models implemented based on Keras.

### 2.5 Integrating the preference of codon composition and the contextual features of TISs

As the neural network in the binary classification scenario mainly discriminates the positive and negative samples based on the contextual sequence features, it may be less sensitive to the difference in the preference of codon composition of TISs. To improve the sensitivity of our method and to make a tradeoff between these two complementary information, here we integrate both *CodonScore* and *ContextScore* to derive the final score representing the likelihood of translation initiation, i.e.,

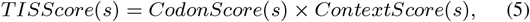

where *s* denotes the sequence profile of a codon site of interest, and a high *T ISScore* is outputted only if both *ContextScore* and *CodonScore* are large enough. Basically, the final score is obtained by weighting between the contributions of the prior preference of codon composition and the sequence contexts to translation initiation.

## 3 Results

### 3.1 TITER accurately predicts TISs

Here, we performed extensive tests to show that TITER greatly outperformed the state-of-the-art methods in predicting TISs, including WRENT (Chew *et al.*, 2016) and PreTIS (Reuter *et al.*, 2016). Note that WRENT cannot compute the initiation scores for those non-AUG codons, while PreTIS only focuses on the 5’ UTR for the near-cognate codons (i.e., the codons differing from AUG by one nucleotide).

After hyperparameter calibration, we first tested our method using a five-fold cross-validation procedure and evaluated its prediction performance based on both the areas under the receiver operating characteristic (AUROC) and the precision recall (AUPR) curves. We found that without using the recurrent layer and the bootstrapping-based technique in our framework, the prediction performance dropped dramatically compared to that of the proposed model (Supplementary Fig. 2), especially for the AUPR score (with drop in AUPR by *∼* 10%), demonstrating the important contributions of these techniques to reduce the modeling variance and boost the prediction accuracy. Also, our model yielded better prediction performance on the AUG codons than non-AUG ones (Supplementary Fig. 2). This indicated that the sequence features of AUG TISs may provide more predictive information in our framework than those of non-AUG TISs. In addition, we observed that even for the AUG sites, our method greatly outperformed WRENT, with increases in AUROC and AUPR scores by 20.9% and 47.0%, respectively.

We also performed additional tests on an independent dataset involving 400 transcripts derived from the Gao15 dataset (see Methods). We first validated the prediction performance of TITER on the Gao15_test dataset, in which the positive and negative samples were selected in the same way as in the construction of our training data, resulting in 935 TISs and 9,098 non-TISs. Test results revealed comparable or even superior prediction performance on this dataset compared to the previous cross-validation results (Figs. 4(a) and 4(b), Supplementary Fig. 2), which further validated that our model did not suffer from the overfitting problem. Similarly, our method also achieved better prediction performance on AUG sites than non-AUG ones, and greatly outperformed WRENT on AUG codons with increases in AUROC and AUPR scores by 17.4% and 46.1%, respectively (Figs. 4(a) and 4(b)). Furthermore, we tested TITER on a more natural and realistic setting, in which we considered all negative samples (i.e., non-TISs) surrounding the positive ones (i.e., TISs) in a transcript. By mainly focusing on those codons with relatively high probability of translation initiation (i.e., *CodonScore >* 1, including AUG, CUG, GUG, UUG and ACG), we selected all the non-TISs of the same triplet before the last TISs for each transcript as negative samples to construct the Gao15_test_extended dataset, which resulted in 767 positive and 9,914 negative samples in total. Test results showed that TITER also yielded excellent prediction performance with AUROC and AURP scores of 89.1% and 61.8%, respectively (Figs. 4(a) and 4(b)). Specifically, the prediction performance was improved after incorporating the prior knowledge on the preference of codon composition of TISs (i.e., *CodonScore*), with an increase of AUPR score by 9.3% (Figs. 4(a) and 4(b)), which thus demonstrated the necessity of our integrative modeling.

**Fig. 4.**
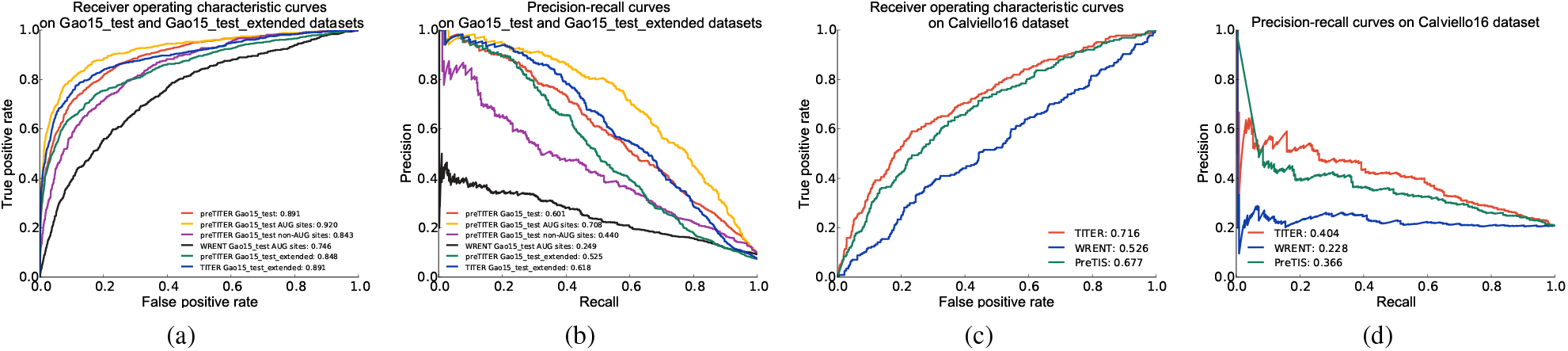
Prediction performance on different test datasets. (a-b) Comparison of prediction performance between different methods on the Gao15 dataset evaluated by (a) ROC and (b) PR curves, respectively. (c-d) Comparison of prediction performance between different methods on the Calviello16 dataset evaluated by (c) ROC and (d) PR curves, respectively. “preTITER” denotes a preliminary version of our deep learning framework that only considered the context features of TISs.

To further validate the generalization of our framework across different datasets and organisms, we additionally evaluated the prediction performance of TITER on an additional dataset of mouse (denoted by Gao15_mouse), which was also derived from Gao *et al.* (2015). All the data preprocess procedures were the same as those for the human data (i.e., the Gao15 dataset). In particular, we randomly selected 360 transcripts as the test data and used the remaining transcripts as our training data. The final prediction performance was evaluated on the test data. Based on the same hyperparameter values calibrated on the Gao15 dataset (Supplementary Table 1), we found that the prediction performance of TITER on the mouse data was comparable or even superior to that on the human data, demonstrating the generalization capacity of our framework (Supplementary Fig. 4).

To facilitate the comparison between TITER and PreTIS, we also constructed a separated uTIS test dataset (denoted by Calviello16). Specifically, this dataset included AUG TISs identified by RiboTaper (Calviello *et al.*, 2016), a statistical method to define the open reading frames (ORFs) through the three-nucleotide periodicity of the ribosome profiling data. The TISs from the transcripts and their isoforms used for training either TITER and PreTIS were excluded, resulting in 383 available transcripts. Furthermore, since the feature engineering of PreTIS required the transcript harboring the TIS of interest to have an orthologous mouse sequence and to be at least 99 nts downstream from its transcript start site, we also excluded all the samples that failed to meet these requirements (*∼*164 transcripts), leaving 253 transcripts available for the test. The positive and negative samples were labeled using the same method as described in (Reuter *et al.*, 2016), resulting in an imbalanced dataset containing 227 positive and 864 negative samples. Tests on the Calviello16 dataset showed that TITER still greatly outperformed both WRENT and PreTIS in this independent dataset, with increases in AUROC by 19.0% and 3.9%, respectively, and in AUPR by 17.6% and 3.8%, respectively (Figs. 4(c) and 4(d)). Note that here the comparison only focused on the AUG codons since RiboTaper only identified the ORFs starting with AUG. In addition, TITER yielded excellent prediction performance on the Gao15 dataset when considering uTISs (still outperforming PreTIS) and dTISs separately (Supplementary Fig. 5).

**Fig. 5.**
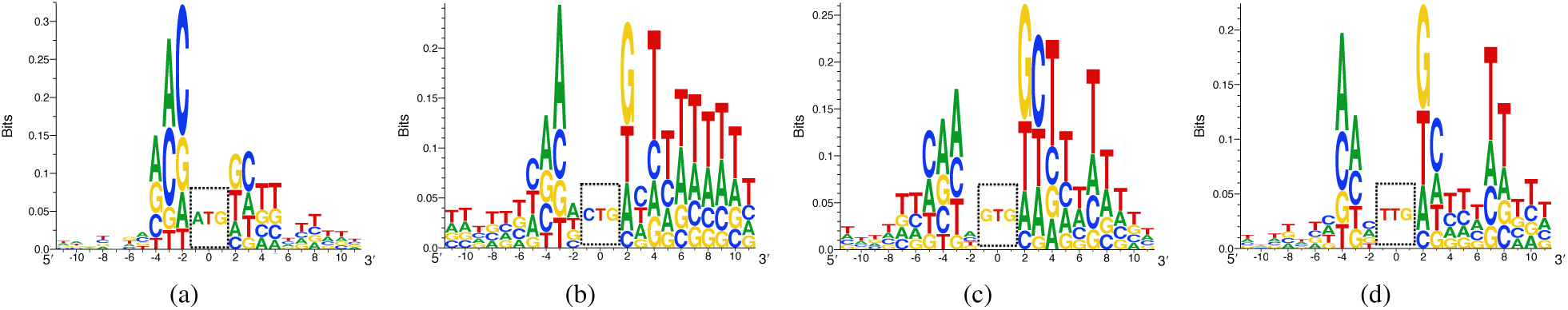
The sequence motifs generated by TITER for (a) AUG (ATG), (b) CUG (CTG), (c) GUG (GTG) and (d) UUG (TTG) TIS codons, respectively. The final position weight matrix (PWM) for each TIS codon was calculated by averaging the optimal input sequences computed by the ensemble of 32 deep neural networks in TITER. The base sequence motifs were visualized using Seq2Logo v2.0 (Thomsen and Nielsen, 2012). All the sequence motifs were visualized in the cDNA setting.

### 3.2 TITER captures the sequence motifs of different TIS codons

The hybrid deep neural network of TITER can be easily extended to generate the sequence motifs of different TIS codons. Based on the similar idea to that used in (Simonyan *et al.*, 2013), here we generated the sequence motifs of a particular TIS codon by optimizing the following problem:

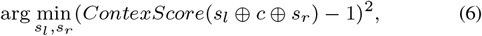

in which *s*_*l*_, *s*_*r*_ and *c* stand for the left sequence, the right sequence and the TIS codon of interest, respectively, and ⊕ represents the sequence concatenation operation. Basically, the above formulation minimizes the difference between the predicted and the positive labels, and finds the optimal sequence with the highest probability of being a TIS for that particular codon. By fixing the weights of a trained neural network, we can optimize the above problem using the gradient descent technique.

Here, we mainly focused on the motif of a region covering both upstream and downstream 10 nts from the TIS codon of interest. In particular, for the canonical AUG start codon, our generated motif was quite similar to the Kozak sequence, i.e., (gcc)gccRccAUGG, which was previously validated to be consensus for aTISs (Kozak, 1989) (Fig. 5(a)). In the Kozak sequence, a lower case indicates a commonly occurred base, a upper case denotes a highly conserved base, and the sequence in a bracket is of uncertain significance. However, simply depending on the Kozak sequence or similar methods like WRENT cannot accurately identify the experimentally-verified AUG TISs (Fig. 4), as the prior knowledge on the sequence motif surrounding the annotated AUG TISs may lead to a biased estimation, which may also explain the slight difference between our generated sequence motif and the Kozak sequence (Fig. 5(a)). Note that the Kozak sequence and the position weight matrix (PWM) based methods (e.g., WRENT) simply assume the independence between different base positions, which largely simplifies the sequence features contributing to translation initiation. In contrast, the deep neural network of TITER can capture more abundant information, such as the correlation between bases in distinct positions, which is generally also relevant to the prediction of translation initiation.

We also generated the sequence motifs for other three NUG TIS codons, including CUG, GUG and UUG. Interestingly, we found that these codons own specific motifs different from that of AUG, especially in the upstream region, indicating that there may exist a different mechanism for alternative translation initiation (Figs. 5(b), 5(c) and 5(d)). Moreover, we observed that these three codons exhibit an (AU)-rich motif in their local downstream regions. As the (AU)-rich regions of mRNAs are commonly associated with high free energy and weak secondary structure (Waterman and Smith, 1978; Lehninger *et al.*, 2008), this implied that the unstructured regions may assist the translation initiation and the following translation elongation process, which can also be supported by known evidence from the previous study (Chew *et al.*, 2016).

### 3.3 Prediction of TITER correlates with translation efficiency

Previous studies have shown that translation initiation at aTISs and uTISs can play important functional roles in regulating gene expression (Ferreira *et al.*, 2013; Chew *et al.*, 2016; Hinnebusch *et al.*, 2016; Calvo *et al.*, 2009). In particular, it has been believed that the strength of translation initiation signals at aTISs can positively correlate with translation efficiency, while the occurrence of uTISs may repress the expression of the main ORFs (Ferreira *et al.*, 2013; Chew *et al.*, 2016; Hinnebusch *et al.*, 2016; Calvo *et al.*, 2009; Jackson *et al.*, 2010; Sonenberg and Hinnebusch, 2009; Hershey *et al.*, 2012). Here, we were particularly interested in investigating the contributions of the predicted *T ISScores* at aTISs and uTISs to the translation efficiency of the main ORFs.

Here, we defined the score of translation efficiency (TE) as the logarithm of the protein expression level divided by the corresponding mRNA expression level. Specifically, the tandem mass spectrometry (MS/MS) data and the mRNA-seq data of HEK293 cell line were obtained from the previously published studies (Geiger *et al.*, 2012; Nam *et al.*, 2014) and were used to derive the levels of protein and mRNA expression, respectively. The iBAQ normalized intensity and the RPKM value were averaged among different replicates for protein and mRNA expression, respectively. We only considered those proteins that were detected in at least two out of three replicates in the MS/MS data. The Uniprot IDs of the genes measured by mass spectrometry were matched to the Ensembl transcript IDs by the Uniprot Retrieve/ID mapping interface (Consortium, 2015). As the current tandem mass spectrum technique cannot accurately distinguish isoforms, our analysis was carried out on the gene level as in Calvo *et al.* (2009). Specifically, for a Uniprot ID that was mapped to multiple transcripts, we selected the transcript with the largest expression value measured by mRNA-seq. The final dataset of matched proteins and mRNAs contained 5,752 genes in total. Consistent with the previous report (Lundberg *et al.*, 2010), we also observed a certain level of correlation between protein and mRNA expression levels (Spearman’s correlation coefficient *r* = 0.67), validating the quality of this dataset to some extent. For uTISs, we followed the same definition of uORF as described in (Calvo *et al.*, 2009), in which a uORF is defined as a continuous segment of codons that has a start codon in the 5’ UTR, a stop codon before the end of the main ORF and a minimum length of nine nts (including the stop codon).

We first employed a linear regression model (implemented based on the scikit-learn library (Pedregosa *et al.*, 2011)) to predict the TE value from the *T ISScore* of aTIS for each transcript. We applied a 10-fold cross-validation procedure to evaluate the correlation between the predicted and the experimentally-derived values. Test results showed that the predicted TE values based on the *T ISScores* of aTISs computed by TITER well correlated with the experimentally-derived TE values (Spearman’s correlation coefficient *r* = 0.234; Table 1), indicating that *T ISScore* can provide predictive power to estimate the strength of translation initiation. Note that there are a variety of biological processes between translation initiation and the final translated protein products that were not included in our modeling, e.g., translation elongation and protein folding, which may explain the weakness of the observed correlation in this test. We further integrated the *T ISScores* of those uTISs that were confidently predicted to form a uORF and with *CodonScore >* 1 and *ContextScore >* 0.95 into the regression model. Interestingly, after integrating the *T ISScores* of uTISs, the correlation between the predicted and experimentally-derived TE values increased from 0.234 to 0.245 (Table 1). In particular, the feature coefficients in the regression model for the *T ISScores* of eligible uTISs were negative, indicating the repressive effect of uORF on the protein expression. Such an increase in correlation was limited, which was consistent with the previous report that the repressive effect of uORF on protein expression level is relatively weak in human (Chew *et al.*, 2016). To further validate the effectiveness of the *T ISScore*, we also performed the same prediction task by replacing the *T ISScores* of uTISs with the number of AUG codons in the 5’ UTR. We found that with this feature replacement, the difference in correlation was almost negligible (Table 1), which thus demonstrated the necessity of considering the contextual sequence features of TISs in the prediction model. We also considered the length of 5’ UTR (which is presumably related to the number of undetected uORFs) in the regression model to test whether those undetected uORFs are influencing TE other than the detected TISs. Test results showed that this additional feature did not affect the prediction performance largely (Table 1), implying that very unlikely our method can suffer from the false negative issue.

### 3.4 TITER quantifies mutational effects on translation initiation

Previously, several studies have identified mutations that are putative to affect translation efficiency through the alteration in the sequence contexts of TISs (Noderer *et al.*, 2014; Kozak, 2002; Wolf *et al.*, 2011) or the introduction of extra uORFs (Hinnebusch *et al.*, 2016; Barbosa *et al.*, 2013; Calvo *et al.*, 2009). As TITER has been shown to be able to accurately predict *bona fide* TISs in human transcriptome, we further explored its ability to quantify the mutational effects that may be related to the physiological and pathological conditions. In particular, we selected two sets of mutations that have been validated through different quantitative reporter assays in (Noderer *et al.*, 2014) and (Calvo *et al.*, 2009), respectively, and then evaluated how the changes of the predicted scores associated with the mutations can reflect their functional effects *in vivo*.

Through flow cytometry, Noderer *et al.* (Noderer *et al.*, 2014) quantitatively measured the effects of seven mutations derived from the COSMIC database (Forbes *et al.*, 2015) and observed consistent effects with other known tumor expression patterns (Fig. 6(a)). Specifically, they employed a dual fluorescence vector with a GFP reporter under the control of a specific TIS context as well as an independent IRES-driven RFP reporter as the internal standard, and the final result was reported with the GFP/RFP ratio. As the experiments were performed by expressing the reporter from a plasmid, we fed the plasmid sequences containing each TIS into the TITER framework, and calculated the changes of the predicted scores (i.e., *ContextScore*) along with the mutations. As expected, the changes of *ContextScores* were in good agreement with the changes of the experimentally-measured GFP/RFP ratios (Fig. 6(b); Pearson’s correlation coefficient *r* = 0.83), indicating that TITER was able to capture the TIS contexts that are related to translation efficiency, even though the mutation information was not included in our training data.

In another study, Calvo *et al.* (Calvo *et al.*, 2009) carried out a series of luciferase assays to demonstrate the effects of the additionally introduced non-overlapping uORFs on protein expression of clotting factor XII (FXII). Their study involved six sequence variants that were associated with the non-overlapping uORFs (Fig. 6(c)). Similar to the above analysis, the sequence contexts of individual TISs, including both aTISs and uTISs, were fed into the TITER framework as input. We observed a good correlation even when only considering the changes of the prediction scores for aTISs (Pearson’s correlation coefficient *r* = 0.75). Notably, the correlation was further improved (Pearson’s correlation coefficient *r* = 0.85), when the *ContextScores* of uTISs were included through a linear leaky scanning model proposed by Ferreira *et al.* (Ferreira *et al.*, 2013) (Fig. 6(d)), i.e.,

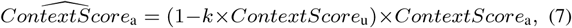

where 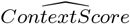 stands for the calibrated *ContextScore*, “a” and “u” denote the “annotated” and “upstream” TISs, respectively, and *k* is the model parameter. Note that here we set *k* = 0.86 based on the previous result of the synthetic reporter assay reported in (Ferreira *et al.*, 2013). Together with our previous genome-wide analysis, these results demonstrated the ability of TITER to quantitatively evaluate the repressive effects of uTIS/uORF on protein expression.

**Fig. 6.**
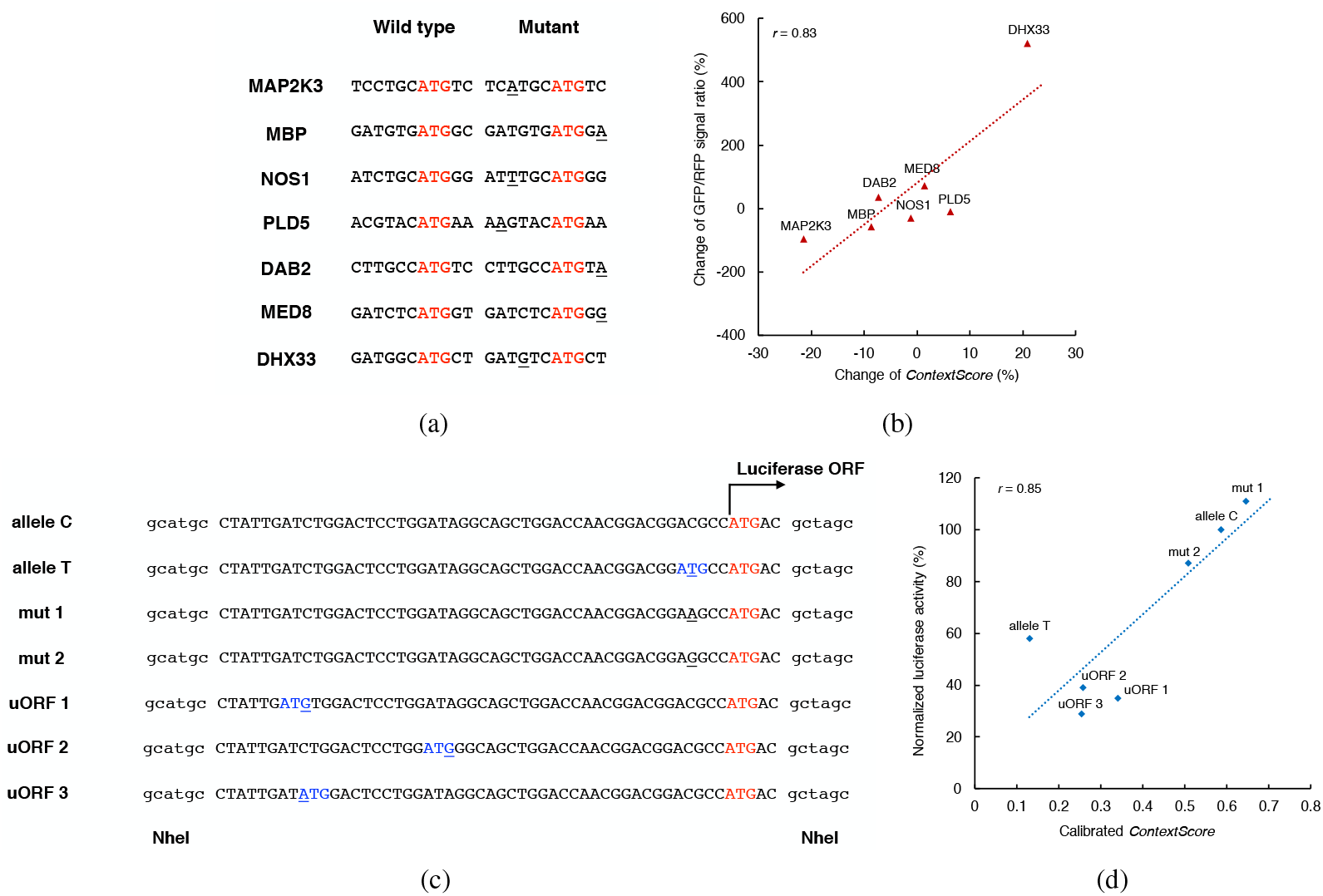
The prediction scores of TITER correlate with the experimentally-measured mutational effects. (a) and (c) Illustrations of different mutations in the tests derived from studies (Noderer et al., 2014) and (Calvo et al., 2009), respectively, in which the base sequences were shown in the cDNA setting. The single nucleotide variants (SNVs) were underlined, and the wild type ATGs and the emerging ATGs were colored in red and blue, respectively. (b) and (d) The correlations between the prediction scores of TITER and the experimentally-measured mutational effects in the previous studies (Noderer et al., 2014) and (Calvo et al., 2009), respectively.

To further confirm that the signal differences presented above were not due to the experimental bias and truly reflected the physiological or pathological effects, we also performed two additional analyses where the input sequences were changed to the corresponding sequences of the real transcripts. As expected, the changes of *ContextScores* for the transcript sequences still maintained a good consistency with the changes of experimental signals in both studies, with only a moderate fluctuation in the correlation, which thus further validated the biological relevance of our results (Supplementary Fig. 3).

## 4 Discussion

The prediction of translation initiation sites (TISs) has long been considered an important task in the studies of gene expression regulation (Jackson *et al.*, 2010; Sonenberg and Hinnebusch, 2009; Hershey *et al.*, 2012). For years, progress has been made on the the identification of the contextual sequence features that can prompt translation initiation, with the focus on the AUG start codon of the main ORFs (Kozak, 1989; Zien *et al.*, 2000; Hatzigeorgiou, 2002; Zur and Tuller, 2013; Chew *et al.*, 2016). However, the existence of alternative initiation codons and multiple initiation positions *in vivo* further complicate this prediction problem. Herein, we have developed a deep learning based framework, called TITER, that can automatically extract intrinsic sequence features from the experimentally identified TISs. By integrating both the preference of codon composition and the contextual sequence information, our unified framework was able to accurately predict various types of TISs. The subsequent motif analysis has expanded our current understanding of the sequence contexts of favored TIS codons. Furthermore, additional analyses of gene expression and mutations showed that TITER can be applied to accurately estimate the probability of translation initiation in various biological scenarios.

Recently, Reuter *et al.* proposed a linear model, called PreTIS, that was trained based on ribosome profiling data and can predict both AUG and non-AUG TISs in the 5’ UTR (Reuter *et al.*, 2016). However, the application of this method is limited by the codon position (i.e., at least 99 nts downstream from the transcript start site and in the 5’ UTR) and the existence of mouse orthologous. Since TITER does not rely on any explicit feature engineering, it possesses the generality of using any input sequence for prediction. Moreover, the extensive tests have shown that TITER can greatly outperform PreTIS, further demonstrating the superiority of our method.

Previously, a number of techniques based on ribosome profiling have been developed to identify and characterize the translation initiation sites. In particular, Lee *et al.* developed the GTI-seq technique, which employed an initiation-specific small molecule ribosome inhibitor, called lactimidomycin (LTM), to capture TISs with both AUG and alternative codons on a genome-wide scale (Lee *et al.*, 2012). As an updated version of this method, Gao *et al.* proposed a dual inhibition technique, called QTI-seq, to address the limitation of the previous method regarding the amplification of ribosome signals on TISs and the inflation of signals at the 5’ ends of transcripts, which achieved a quantitative profiling of initiating ribosomes (Gao *et al.*, 2015). Moreover, they also applied stringent mapping protocol and statistical test to increase the fidelity of the identified TISs (Gao *et al.*, 2015). Therefore, in our study we mainly chose the dataset generated from (Gao *et al.*, 2015) to facilitate the training and test of TITER. With the careful selection of data source and the accurate prediction performance on various test settings, we believe that TITER will be useful for the community to investigate probable TISs and further expand our understanding of the mechanisms underlying translation initiation.

**Funding**

This work was supported in part by the National Basic Research Program of China Grant 2011CBA00300 and 2011CBA00301, the National Natural Science Foundation of China Grant 61033001, 61361136003 and 61472205, the US National Science Foundation Grant DBI-1262107 and IIS-1646333, the China’s Youth 1000-Talent Program, and the Beijing Advanced Innovation Center for Structural Biology.

## Footnotes

1 https://keras.io/

2 https://github.com/maxpumperla/hyperas/

## References

Aken, B. L., Ayling, S., Barrell, D., Clarke, L., Curwen, V., Fairley, S., Fernandez Banet, J., Billis, K., García Girón, C., Hourlier, T., Howe, K., Kähäri, A., Kokocinski, F., Martin, F. J., Murphy, D. N., Nag, R., Ruffier, M., Schuster, M., Tang, Y. A., Vogel, J.-H., White, S., Zadissa, A., Flicek, P., and Searle, S. M. J. (2016). The Ensembl gene annotation system. Database, 2016.

Alipanahi, B., Delong, A., Weirauch, M. T., and Frey, B. J. (2015). Predicting the sequence specificities of DNA-and RNA-binding proteins by deep learning. Nat Biotech, 33(8), 831–838.

Barbosa, C., Peixeiro, I., and Romão, L. (2013). Gene expression regulation by upstream open reading frames and human disease. PLOS Genetics, 9(8), e1003529–.

Bengio, Y. (2012). Neural Networks: Tricks of the Trade: Second Edition, chapter Practical Recommendations for Gradient-Based Training of Deep Architectures, pages 437–478. Springer Berlin Heidelberg, Berlin, Heidelberg.

Bergstra, J. S., Bardenet, R., Bengio, Y., and Kégl, B. (2011). Algorithms for hyper-parameter optimization. In J. Shawe-Taylor, R. S. Zemel, P. L. Bartlett, F. Pereira, and K. Q. Weinberger, editors, Advances in Neural Information Processing Systems 24, pages 2546–2554. Curran Associates, Inc.

Calviello, L., Mukherjee, N., Wyler, E., Zauber, H., Hirsekorn, A., Selbach, M., Landthaler, M., Obermayer, B., and Ohler, U. (2016). Detecting actively translated open reading frames in ribosome profiling data. Nat Meth, 13(2), 165–170.

Calvo, S. E., Pagliarini, D. J., and Mootha, V. K. (2009). Upstream open reading frames cause widespread reduction of protein expression and are polymorphic among humans. Proceedings of the National Academy of Sciences, 106(18), 7507–7512.

Chew, G.-L., Pauli, A., and Schier, A. F. (2016). Conservation of uORF repressiveness and sequence features in mouse, human and zebrafish. Nature Communications, 7, 11663–.

Collobert, R., Weston, J., Bottou, L., Karlen, M., Kavukcuoglu, K., and Kuksa, P. (2011). Natural language processing (almost) from scratch. J. Mach. Learn. Res., 12, 2493–2537.

Consortium, T. U. (2015). Uniprot: A hub for protein information. Nucleic Acids Research, 43(D1), D204–D212.

Denil, M., Bazzani, L., Larochelle, H., and de Freitas, N. (2012). Learning where to attend with deep architectures for image tracking. Neural Computation, 24(8), 2151–2184.

Ferreira, J. P., Overton, K. W., and Wang, C. L. (2013). Tuning gene expression with synthetic upstream open reading frames. Proceedings of the National Academy of Sciences, 110(28), 11284–11289.

Forbes, S. A., Beare, D., Gunasekaran, P., Leung, K., Bindal, N., Boutselakis, H., Ding, M., Bamford, S., Cole, C., Ward, S., Kok, C. Y., Jia, M., De, T., Teague, J. W., Stratton, M. R., McDermott, U., and Campbell, P. J. (2015). COSMIC: Exploring the world's knowledge of somatic mutations in human cancer. Nucleic Acids Research, 43(D1), D805–D811.

Gao, X., Wan, J., Liu, B., Ma, M., Shen, B., and Qian, S.-B. (2015). Quantitative profiling of initiating ribosomes *in vivo*. Nat Meth, 12(2), 147–153.

Geiger, T., Wehner, A., Schaab, C., Cox, J., and Mann, M. (2012). Comparative proteomic analysis of eleven common cell lines reveals ubiquitous but varying expression of most proteins. Molecular & Cellular Proteomics, 11(3).

Hassanzadeh, H. R. and Wang, M. D. (2016). DeeperBind: Enhancing prediction of sequence specificities of DNA binding proteins. In IEEE International Conference on Bioinformatics and Biomedicine, BIBM 2016, Shenzhen, China, December 15-18, 2016, pages 178–183.

Hatzigeorgiou, A. G. (2002). Translation initiation start prediction in human cDNAs with high accuracy. Bioinformatics, 18(2), 343–350.

Hershey, J. W., Sonenberg, N., and Mathews, M. B. (2012). Principles of translational control: An overview. Cold Spring Harbor Perspectives in Biology, 4(12).

Hinnebusch, A. G., Ivanov, I. P., and Sonenberg, N. (2016). Translational control by 5'-untranslated regions of eukaryotic mRNAs. Science, 352(6292), 1413–1416.

Hinton, G., Deng, L., Yu, D., Dahl, G. E., Mohamed, A.-r., Jaitly, N., Senior, A., Vanhoucke, V., Nguyen, P., Sainath, T. N., and Kingsbury, B. (2012). Deep neural networks for acoustic modeling in speech recognition: The shared views of four research groups. Signal Processing Magazine, IEEE, 29(6), 82–97.

Hinton, G. E. and Salakhutdinov, R. R. (2006). Reducing the dimensionality of data with neural networks. Science, 313(5786), 504–507.

Hinton, G. E., Osindero, S., and Teh, Y.-W. (2006). A fast learning algorithm for deep belief nets. Neural Comput., 18(7), 1527–1554.

Ingolia, N. T., Ghaemmaghami, S., Newman, J. R. S., and Weissman, J. S. (2009). Genome-wide analysis in vivo of translation with nucleotide resolution using ribosome profiling. Science, 324(5924), 218–223.

Ingolia, N. T., Brar, G. A., Rouskin, S., McGeachy, A. M., and Weissman, J. S. (2012). The ribosome profiling strategy for monitoring translation *in vivo* by deep sequencing of ribosome-protected mRNA fragments. Nat. Protocols, 7(8), 1534–1550.

Jackson, R. J., Hellen, C. U. T., and Pestova, T. V. (2010). The mechanism of eukaryotic translation initiation and principles of its regulation. Nat Rev Mol Cell Biol, 11(2), 113–127.

Kozak, M. (1989). Context effects and inefficient initiation at non-AUG codons in eucaryotic cell-free translation systems. Molecular and Cellular Biology, 9(11), 5073–5080.

Kozak, M. (2002). Emerging links between initiation of translation and human diseases. Mammalian Genome, 13(8), 401–410.

Larochelle, H. and Hinton, G. (2010). Learning to combine foveal glimpses with a third-order boltzmann machine. In J. Lafferty, C. Williams, J. Shawe-Taylor, R. Zemel, and A. Culotta, editors, Advances in Neural Information Processing Systems 23, pages 1243–1251.

Lee, S., Liu, B., Lee, S., Huang, S.-X., Shen, B., and Qian, S.-B. (2012). Global mapping of translation initiation sites in mammalian cells at single-nucleotide resolution. Proceedings of the National Academy of Sciences, 109(37), E2424–E2432.

Lehninger, A., Nelson, D., and Cox, M. (2008). Lehninger Principles of Biochemistry.

W. H. Freeman. Li, H. and Jiang, T. (2005). A class of edit kernels for SVMs to predict translation initiation sites in eukaryotic mRNAs. Journal of Computational Biology, 12(6), 702–718.

Lundberg, E., Fagerberg, L., Klevebring, D., Matic, I., Geiger, T., Cox, J., Algenäs, C., Lundeberg, J., Mann, M., and Uhlen, M. (2010). Defining the transcriptome and proteome in three functionally different human cell lines. Molecular Systems Biology, 6(1).

Michel, A. M., Andreev, D. E., and Baranov, P. V. (2014). Computational approach for calculating the probability of eukaryotic translation initiation from ribo-seq data that takes into account leaky scanning. BMC Bioinformatics, 15(1), 380.

Nam, J.-W., Rissland, O. S., Koppstein, D., Abreu-Goodger, C., Jan, C. H., Agarwal, V., Yildirim, M. A., Rodriguez, A., and Bartel, D. P. (2014). Global analyses of the effect of different cellular contexts on MicroRNA targeting. Molecular Cell, 53(6), 1031–1043.

Noderer, W. L., Flockhart, R. J., Bhaduri, A., Diaz de Arce, A. J., Zhang, J., Khavari, P. A., and Wang, C. L. (2014). Quantitative analysis of mammalian translation initiation sites by FACS-seq. Mol Syst Biol, 10(8), 748–.

Peabody, D. S. (1989). Translation initiation at non-AUG triplets in mammalian cells. Journal of Biological Chemistry, 264(9), 5031–5.

Pedregosa, F., Varoquaux, G., Gramfort, A., Michel, V., Thirion, B., Grisel, O., Blondel, M., Prettenhofer, P., Weiss, R., Dubourg, V., Vanderplas, J., Passos, A., Cournapeau, D., Brucher, M., Perrot, M., and Duchesnay, E. (2011). Scikitlearn: Machine learning in Python. Journal of Machine Learning Research, 12, 2825–2830.

Quang, D. and Xie, X. (2016). DanQ: A hybrid convolutional and recurrent deep neural network for quantifying the function of DNA sequences. Nucleic Acids Research, 44(11), e107.

Reuter, K., Biehl, A., Koch, L., and Helms, V. (2016). PreTIS: A tool to predict non-canonical 5' UTR translational initiation sites in human and mouse. PLOS Computational Biology, 12(10), e1005170–.

Rumelhart, D. E., Hinton, G. E., and Williams, R. J. (1986). Learning representations by back-propagating errors. Nature, 323(6088), 533–536.

Simonyan, K., Vedaldi, A., and Zisserman, A. (2013). Deep inside convolutional networks: Visualising image classification models and saliency maps. CoRR, abs/1312.6034.

Sonenberg, N. and Hinnebusch, A. G. (2009). Regulation of translation initiation in eukaryotes: Mechanisms and biological targets. Cell, 136(4), 731–745.

Srebro, N., Rennie, J., and Jaakola, T. (2005). Maximum-margin matrix factorization. In Advances in Neural Information Processing Systems 17, volume 17, pages 1329–1336.

Srivastava, N., Hinton, G., Krizhevsky, A., Sutskever, I., and Salakhutdinov, R. (2014). Dropout: A simple way to prevent neural networks from overfitting. J. Mach. Learn. Res., 15(1), 1929–1958.

Thomsen, M. C. F. and Nielsen, M. (2012). Seq2Logo: A method for construction and visualization of amino acid binding motifs and sequence profiles including sequence weighting, pseudo counts and two-sided representation of amino acid enrichment and depletion. Nucleic Acids Research, 40(W1), W281–W287.

Wallace, B., Small, K., Brodley, C., and Trikalinos, T. (2011). Class imbalance, redux. In 2011 IEEE 11th International Conference on Data Mining, pages 754–763.

Waterman, M. and Smith, T. (1978). RNA secondary structure: A complete mathematical analysis. Mathematical Biosciences, 42(3), 257–266.

Wolf, A., Caliebe, A., Thomas, N. S., Ball, E. V., Mort, M., Stenson, P. D., Krawczak, M., and Cooper, D. N. (2011). Single base-pair substitutions at the translation initiation sites of human genes as a cause of inherited disease. Hum. Mutat., 32(10), 1137–1143.

Zhang, S., Zhou, J., Hu, H., Gong, H., Chen, L., Cheng, C., and Zeng, J. (2015). A deep learning framework for modeling structural features of RNA-binding protein targets. Nucleic Acids Research, 44(4), e32.

Zhang, S., Hu, H., Zhou, J., He, X., Jiang, T., and Zeng, J. (2016). ROSE: A deep learning based framework for predicting ribosome stalling. bioRxiv.

Zhou, J. and Troyanskaya, O. G. (2015). Predicting effects of noncoding variants with deep learning-based sequence model. Nat Meth, 12(10), 931–934.

Zien, A., Rätsch, G., Mika, S., Schölkopf, B., Lengauer, T., and Müller, K.-R. (2000). Engineering support vector machine kernels that recognize translation initiation sites. Bioinformatics, 16(9), 799–807.

Zur, H. and Tuller, T. (2013). New universal rules of eukaryotic translation initiation fidelity. PLOS Computational Biology, 9(7), e1003136–.

